# mRNA-LNP COVID-19 vaccine lipids induce low level complement activation and production of proinflammatory cytokines: Mechanisms, effects of complement inhibitors, and relevance to adverse reactions

**DOI:** 10.1101/2024.01.12.575122

**Authors:** Tamás Bakos, Tamás Mészáros, Gergely Tibor Kozma, Petra Berényi, Réka Facskó, Henriette Farkas, László Dézsi, Carlo Heirman, Stefaan de Koker, Raymond Schiffelers, Kathryn Anne Glatter, Tamás Radovits, Gábor Szénási, János Szebeni

## Abstract

Messenger RNA-containing lipid nanoparticles (mRNA-LNPs) enabled widespread COVID-19 vaccination with a small fraction of vaccine recipients displaying acute or sub-acute inflammatory symptoms. The molecular mechanism of these adverse events (AEs) remains undetermined. Here we report that the mRNA-LNP vaccine, Comirnaty, triggers low-level complement (C) activation and production of inflammatory cytokines, which may be key underlying processes of inflammatory AEs. In serum, Comirnaty and the control PEGylated liposome (Doxebo) caused different rises of C split products, C5a, sC5b-9, Bb and C4d, indicating stimulation of the classical pathway of C activation mainly by the liposomes, while a stronger stimulation of the alternative pathway was equal with the vaccine and the liposomes. Spikevax had similar C activation as Comirnaty, but viral or synthetic mRNAs had no such effect. In autologous serum-supplemented peripheral blood mononuclear cell (PBMC) cultures, Comirnaty caused increases in the levels of sC5b-9 and proinflammatory cytokines in the following order: IL-1α < IFN-γ < IL-1β < TNF-α < IL-6 < IL-8, whereas heat-inactivation of serum prevented the rises of IL-1α, IL-1β, and TNF-α. Clinical C inhibitors, Soliris and Berinert, suppressed vaccine-induced C activation in serum but did not affect cytokine production when applied individually. These findings suggest that the PEGylated lipid coating of mRNA-LNP nanoparticles can trigger C activation mainly via the alternative pathway, which may be causally related to the induction of some, but not all inflammatory cytokines. While innate immune stimulation is essential for the vaccine’s efficacy, concurrent production of C- and PBMC-derived inflammatory mediators may contribute to some of the AEs. Pharmacological attenuation of harmful cytokine production using C inhibitors likely requires blocking the C cascade at multiple points.

## INTRODUCTION

The nucleoside-modified mRNA COVID-19 vaccines, BNT162b2 (Comirnaty) from Pfizer-BioNTech and mRNA-1273 (Spikevax), from Moderna, are the most widely applied vaccines against SARS-CoV-2 in the United States and many other countries worldwide. These vaccines are considered effective and safe^1–4^, nevertheless, like any medical intervention, a small fraction of vaccine recipients develop acute or subacute side effects, referred to as adverse events (AEs). Some AEs are severe (SAEs), leading to emergency care, hospitalization, permanent disability or even death.

Based mainly on COVID-19 symptoms, an international team of established experts (Brighton Collaboration’s Safety Platform for Emergency Vaccines) assembled a list of COVID-19 vaccine-specific SAEs^5^, whose occurrence was analyzed^6^ with focus on Pfizer-BioNTech’s phase 2/3 trials evaluating the safety, immunogenicity, and efficacy of Comirnaty^7^. This study demonstrated a 36 % higher risk of SAEs in the vaccine group, with 97% of SAEs overlapping with COVID-19 acute and chronic symptoms. These findings suggest that the mRNA-LNP vaccines and the SARS-CoV-2 virus, which share many structural features of lipid membrane-enveloped mRNA nanoparticles, have overlapping harmful impacts on the body. In other words, beyond its mission to code for S-protein (SP) expression by antigen-presenting immune cells, the mRNA-LNP vaccines resemble the virus in causing certain pathologic effects.

Table 1 lists the vaccine’s reported AEs categorized according to the affected organ systems. Of note, myocarditis/pericarditis^8–15^, HSRs/anaphylaxis^16–19^, autoimmune diseases^20^, thrombosis, thrombocytopenia and other coagulation disorders^21–24^ skin^25–28^ and ocular inflammations^29–31^, Guillain-Barre syndrome^32–38^ and other neurologic problems^39, 40^ can all be linked to acute or subacute inflammatory processes that are also characteristic of SARS-CoV-2 infections.

**Table 1.**
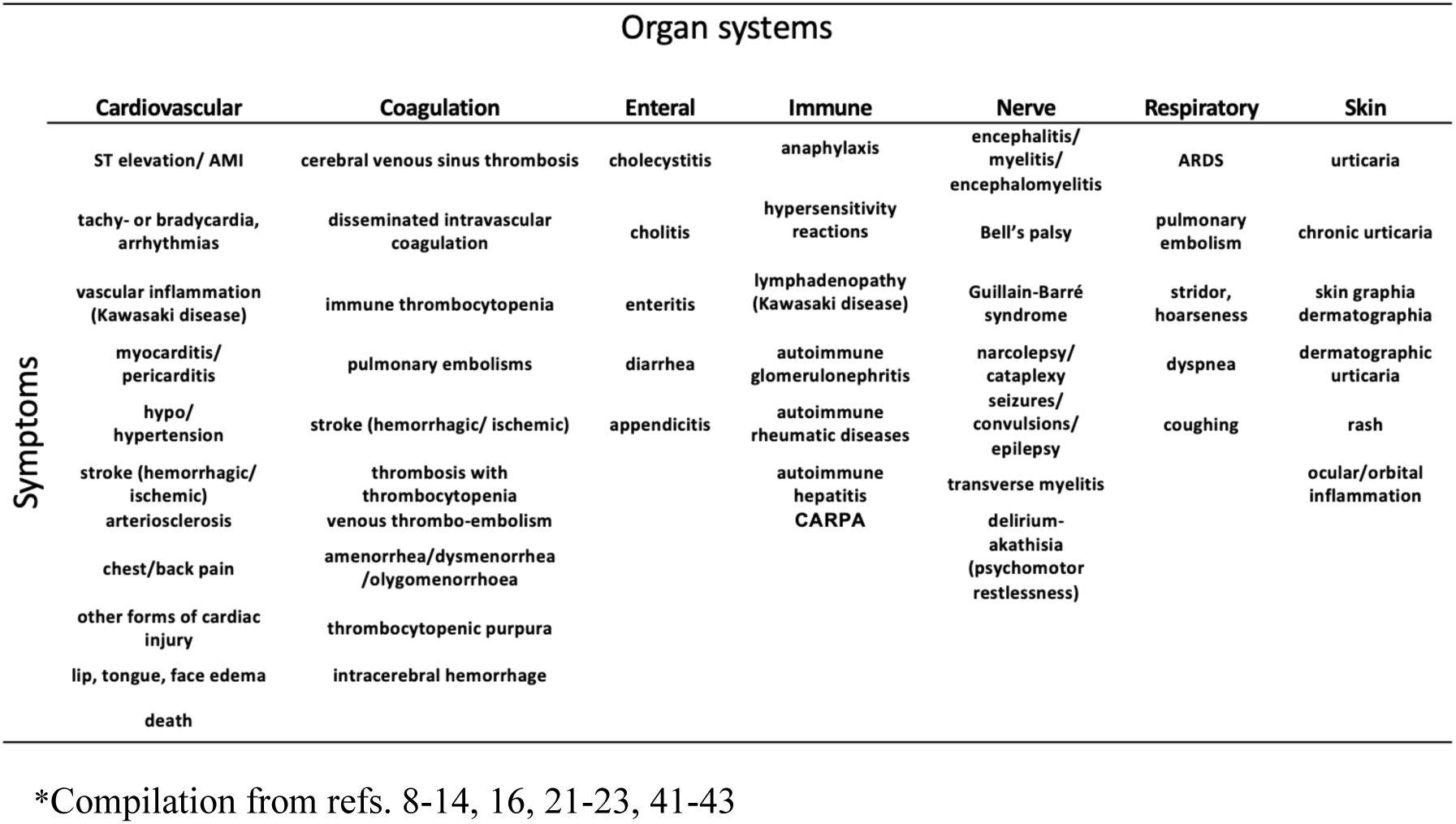
Adverse effects of mRNA-LNP vaccines*.

This coincidence of virus and vaccine-induced inflammatory symptoms points to a common underlying mechanism, and the goal of the present study was to find this common denominator. We focused on two well-known contributors to inflammation: complement (C) activation with production of anaphylatoxins and the release of proinflammatory cytokines. As test agents, we used Pfizer-BioNTech’s Comirnaty, Moderna’s Spikevax, PEGylated liposomes (Doxebo)^44^, naked mRNAs (coding for SARS-CoV-2 and 3 other proteins) and zymosan^45^, as the positive control. The two test systems were human peripheral blood mononuclear cell cultures (PBMC) supplemented with 20% native or heat-inactivated autologous serum^46^.

Our findings reveal that mRNA-LNP nanoparticles (NPs) can activate C and simultaneously induce certain proinflammatory cytokines; phenomena that are also characteristic of severe COVID-19^47–54^. The relationships found between C activation and cytokine release may help in better understanding the mechanism of inflammatory side effects of mRNA-LNP-based vaccines, and in developing new strategies for the prevention not only of vaccine-induced SAEs but also those caused by other medications utilizing the mRNA-LNP technology platform.

## METHODS

### Materials

Mammalian cell culture medium (RPMI-1640, R5), non-essential amino acid solution (0,1 mM), pyruvate solution (1 mM), penicillin-streptomycin antibiotics solution and β-mercaptoethanol (50 µM) Ficoll-Paque, ethylenediaminetetraacetic acid (EDTA), Dulbecco’s phosphate buffered saline (D-PBS) and trypan blue dye were also purchased for Merck Life Science Ltd. (Budapest, Hungary). R5 medium was from ThermoFisher Scientific. Comirnaty and Spikevax vaccines used against the delta variant of SARS-CoV-2 virus (lot numbers: ET7205 and FA 5829) were obtained from Semmelweis University’s Pharmacy. They were stored as instructed by the manufacturers and immediately used after opening the vials within the expiration date. The C inhibitor anti-C5 monoclonal antibody (mAb), Eculizumab (Soliris^®^), was from Alexion Pharmaceuticals (Boston, MA), today owned by Astra Zeneca (Cambridge, UK). The C1-esterase inhibitor (Berinert^®^) was from CSL Behring GmbH (Marburg, Germany), and 5-methoxyuridine-modified SARS-CoV-2 mRNA (E484K, N501Y) was from OZ Biosciences SAS (Marseille, France).

### Ethical permission, donors, and blood collection

The Scientific and Research Ethics Committee of the Medical Research Council of Hungary granted ethical approval for this research project (TUKEB 15576/2018/EKU). After getting informed consent from all adult, healthy participants in this study, their blood was withdrawn by a phlebotomist.

### Separation of serum and PBMC

The serum was separated from whole blood after coagulating the blood (approx. 25 min) at room temperature in a Greiner VACUETTE Z Serum Sep Clot Activator (8 ml) blood collector tube. The tubes were centrifuged (2000 g, 4°C, 15 min), and the supernatants (serum) were collected. PBMCs were separated from uncoagulated whole blood collected in Greiner VACUETTE K2E K_2_EDTA (9 ml) blood collector tubes and diluted in a 1:1 ratio with 6 mM EDTA containing D-PBS using Ficoll Paque density gradient centrifugation (500 g, 25°C, 30 min). Cells that formed a white-colored ring were then suspended in 6 mM EDTA containing D-PBS to ensure the removal of the remaining substances from the previous steps and were centrifuged again (500 g, 25°C, 30 min). To remove thrombocytes and residual EDTA, cells were washed with cold, EDTA-free D-PBS and were centrifuged again (500 g, 4°C, 15 min). Following the centrifugation, the cells were divided into three portions and were suspended in 3 culture media: 1) medium only (R5); 2) R5 containing 20% autologous human serum, representing the in vivo conditions with 1/5 amount of C proteins and other serum components exposed to the cells; 3) R5 containing 20% autologous human serum inactivated by heat, representing the in vivo conditions without activable C proteins. To inactivate sera, also known as decomplementation^55^, a portion of the innate serum from each donor was incubated at 56°C for 30 minutes. The 20% serum ratio was chosen on the basis of preliminary studies showing no significant cell toxicity over 18h, as determined by trypan blue exclusion^46^. The suspensions were centrifuged (500 g, 4°C, 15 min) and subsequently re-suspended in the culture media specified above.

### Preparation of mRNAs

To investigate the C activating effects of naked mRNAs, we applied N1-Methyl Pseudouridine-modified, 5’ cap1 cleancapped mRNAs prepared by Etherna Biopharmaceuticals (Niel, Belgium), as described earlier^56^. The samples coding for the proteins CD40L (29.3 kDa), Cre-recombinase (38.5 kDa), and luciferase (60.7 kDa) contained 1,200, 1,450 and 2,086 nt, respectively. Their sequence and other details are shown the Supplemental data section.

### PBMC Studies

Following separation, the PBMCs were suspended in R5 medium supplemented with 20% native or heat-inactivated autologous serum. The suspension was then placed into the inner wells of a 96-well microtiter plates, with each well containing approximately 5 × 10^5^ cells. The plates were placed in a CO_2_ incubator (5% CO_2_) at 37°C. Aliquots were collected at 45 minutes and 18 hours after the start of the experiment. Additionally, baseline samples (“0 min samples”) were collected before the incubation. After collection, samples were centrifuged (2500 g, 4°C, 10 min), and the supernatants were stored at –80°C until the C and cytokine assays were performed. Soluble terminal C complex (sTCC, sC5b9) was measured with an ELISA kit from Svar Life Science AB (Malmö, Sweden), and for the measurement of cytokines, we used the Q-View™ LS chemiluminescent imager paired with Q-View™ Software, obtained from Quansys Biosciences (West Logan, UT, USA). The cytokine panel supplied by the company was the Human Cytokine Inflammation Panel 1 multiplex ELISA, measuring IL-1α, IL-1β, IL-2, IL-4, IL-6, IL-8, IL-10, IFN-γ, and TNF-α. All procedures followed the manufacturer’s instructions.

### Serum Studies

The vaccine and other test samples were mixed with native serum at a 1:3 volume ratio, followed by incubation at 37°C for 30 or 45 min, as stated in the text. The incubation was stopped by dilution with the kit’s sample diluent supplemented with 10 mM EDTA. Soluble TCC, C5a, C4d and Bb were measured with Svar’s C5b-9 kit (Svar Life Sci., Malmö, Sveden) and TECO*medical*’s C5a, C4d and Bb ELISA kits (TECO*medical* AG, Sissach, Switzerland).

### Statistical analysis

All data reported are means ± SEM. Statistical comparisons were made using paired t-tests or one-way ANOVA and Dunnetts’ or Tukey’s multiple comparisons post hoc tests, as specified in the text.

## RESULTS

### Complement activation by Comirnaty in PBMC cultures

**Figure 1A** shows the levels of soluble terminal complex (sC5b-9) in PBMC supernatants as endpoint of C activation after incubation with Comirnaty. In native serum-supplemented PBMC cultures (solid-colored bars), the averaged data from 5 different blood donors showed significant rise of sC5b-9 at 45 min relative to 0 min baseline both in the absence and the presence of Comirnaty, the rise being more pronounced in the presence of the vaccine. These data suggest that incubation at 37°C for 45 min induces significant spontaneous C activation in PBMC, upon which the vaccine-induced small activation superimposes. Surprisingly, we recorded significant increases in sC5b-9 also in heat-inactivated serum even at baseline, upon which the vaccine caused additional C activation relative to the vaccine-free 45 min samples. This counterintuitive observation is likely related to liberation of heat-stable sC5b-9 upon heat treatment, which is superimposed on spontaneous C activation during incubation. This phenomenon obscured the difference between heated and native serum upon spontaneous C activation, but the Comirnaty-induced small, but significant increase of sC5b-9 was still present. The latter data raise the possibility of incomplete decomplementation upon heat inactivation of serum, or C-independent activation of C5^57^.

**Figure 1.**
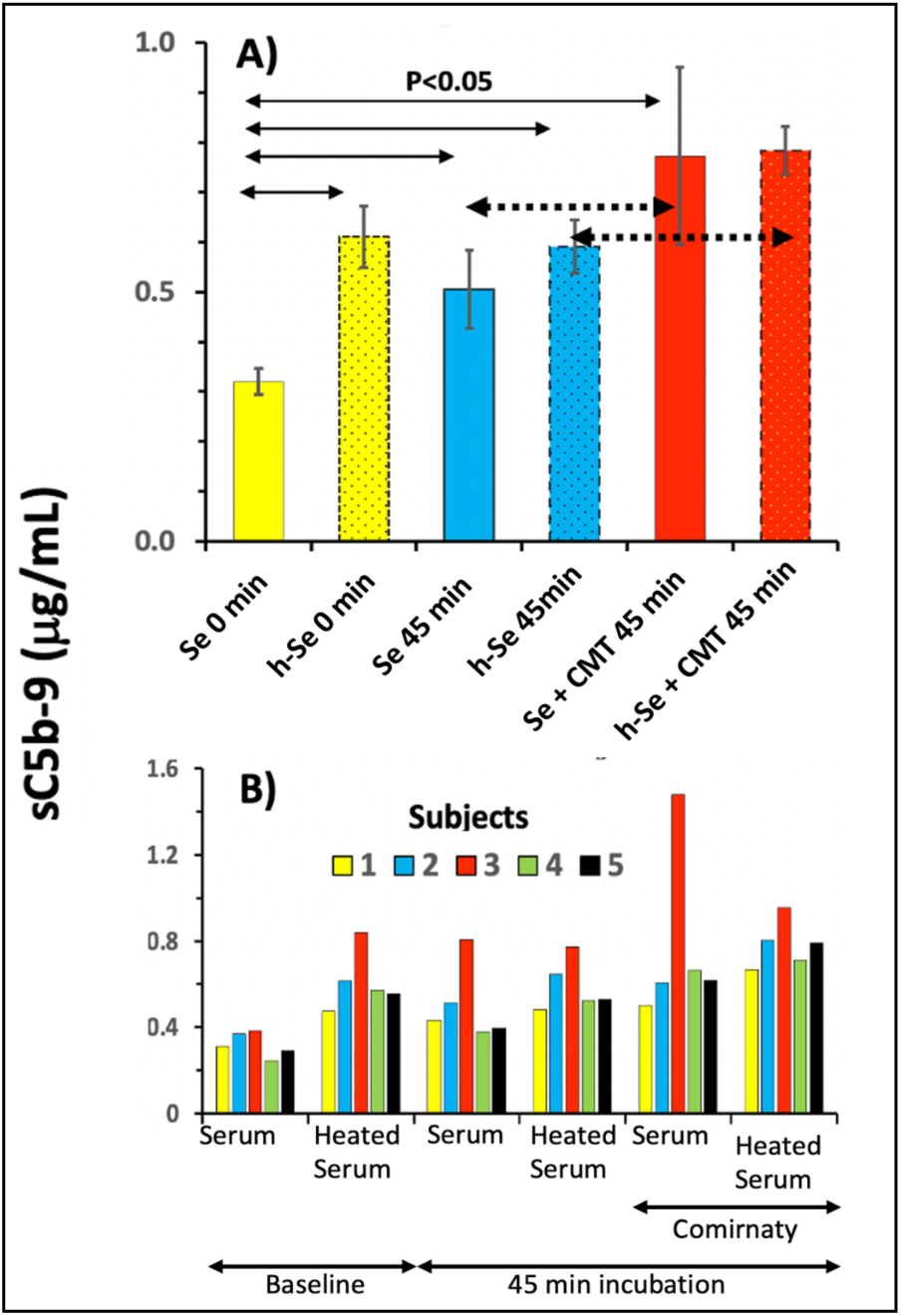
Complement activation by Comirnaty in cultured R5 medium supplemented with 20% native (solid colored bars) or heat-inactivated autologous serum (dash-framed dotted bars). The vaccine was added at 4.8 μg/mL, bars in (A) show averages (± SEM) from 5 donors, yellow, blue and red colors indicate baseline and 45 min incubation in the absence and presence of Comirnaty, respectively. Panel B shows the same data on individual level. The incubation in a CO_2_ incubator at 37°C lasted for 18 h, while aliquots for the sC5b-9 ELISA were taken only after 45 min. The arrows indicate the significance of differences, dotted arrows pair the bars which show vaccine induced C activation either in native or heat-inactivated serum supplemented cultures. Se, serum; h-Se, heated serum, 0 min: before incubation, after adding the serum; 45 min, after 45 min incubation; CMT, Comirnaty

**Figure 1B** individually visualizes the same Comirnaty-induced sC5b-9 changes that were averaged in Panel A. This presentation points to biological, rather than technical cause of variation, since one of the 5 sera (Subject #3) displayed consistently higher (up to 2-fold) sC5b-9 values under all test conditions compared to the other 4 sera. This donor might have had increased sensitivity towards C activation in general, exemplifying the common experience of substantial individual variation in this immune response.

Taken together, these findings in 20% human serum confirm the previous data in 60% pig serum that Comirnaty can activate the C system^17^. Nevertheless, the dilution of serum resulted in a 5-fold reduction of the assay’s sensitivity, thus reducing the effective dynamic range of sC5b-9 changes. This led us to test cell-free native serum which was diluted only by the added test agents by 25%. As shown below, this switch from 20 to 75% serum allowed us to validate the sC5b-9 assay for the quantification of vaccine-induced C activation, to assess its pathways and relative strength, as well as the effects of C inhibitors.

### Features of C activation by Comirnaty in 75% human serum

**Figures 2A-C** show the effects of the mRNA vaccine, Comirnaty (red bar), a PEGylated liposome, Doxebo (blue bar) and zymosan (black bar) on C activation in 75% serum of 5 blood donors, along with the time-matched PBS (baseline) values (yellow bars). To quantitate C activation, we measured concurrent changes in four reaction biomarkers; sC5b-9, a stable end product of the C cascade (**Panel A**); the anaphylatoxin C5a, one of the strongest proinflammatory mediator in blood (**B**); C4d, a C activation biomarker specific to classical and/or lectin pathway activations **(C),** and Bb, a biomarker specific to the alternative pathway C activation (**D**).

**Figure 2.**
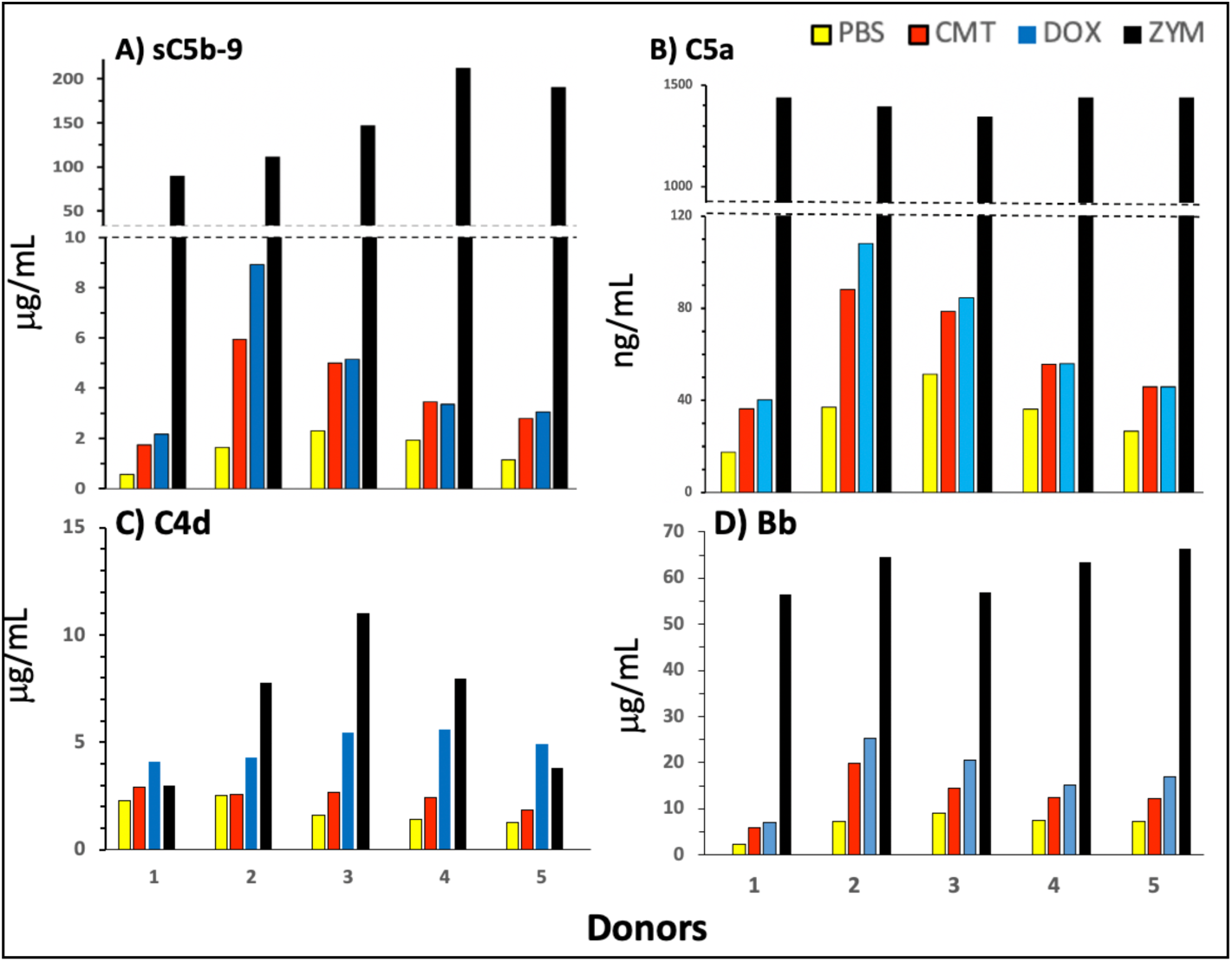
Complement activation by Comirnaty (CMT), control PEGylated liposomes (Doxebo, DOX), and zymosan (ZYM). The total lipid in the Comirnaty and Doxebo samples were 0.64 and 3.99 mg/mL, respectively (STable 1), and the mRNA concentration in Comirnaty was 25 μg/mL. These lipid contents were based on equal, 4-fold dilution of vaccine and liposome stocks in serum, and imply 6.2-fold higher lipid concentrations in Doxebo vs. Comirnaty. All other conditions of the experiments were the same as in Figure 1. Bars with different colors represent different reaction triggers, as defined in the key. The individual rises relative to the respective baselines (n=5) were significant for all 4 reaction markers (A-D) by paired 2 sample t-test at P< 0.05 or <0.01. The MWs of sC5b-9, C5a, C4d and Bb, used for molar concentration calculations, are 1,030.0, 10.4, 47 and 60 kDa, respectively.

Figure 2 shows that both Comirnaty and Doxebo caused small but significant, 50-220% increases in sC5b9 (Figure 2A), C5a (B) and Bb (D) compared to PBS baseline. In contrast, C4d was significantly raised only by Doxebo, the Comirnaty-induced minor rise of C4d was seen only in 3 of the 5 sera. A closer analysis of these data also reveals that the Comirnaty-induced rise of sC5b-9 in different serum donors on the μg mL^−1^ scale (A) was proportional to the rise of C5a on the ng mL^−1^ scale (B). Furthermore, the ∼4 μg/mL and ∼60 ng/mL average rises of sC5b-9 and C5a translates to 4 and 6 nM, respectively (see MWs in the legend), which ratio is a realistic divergence from equimolar formation of sC5b-9 and C5a as only a part of *de novo* formed C5b-9 binds to S-protein (vitronectin) in serum^58–60^ which is measured by the sC5b-9 kit. The above facts, taken together with the proportionality of % sC5b-9 rises and serum % values in Figure 1 and 2A (i.e. ∼4-fold higher sC5b-9 in 75% serum than that in 20% serum in PBMC), also validates the sC5b-9 assay for the quantification of C activation under different conditions and the extrapolation from serum data to immune cell-containing media, such as blood.

Further notable observations in Figure 2 are that Comirnaty and Doxebo caused far lower C activation, <1% of the activity of zymosan (Table 2), and that the Comirnaty and Doxebo-induced rises in sC5b9, C5a and Bb were near equal (Figures 2A, B and D), although, importantly, Comirnaty had ∼7-fold less lipid in the samples than Doxebo (see legend).

**Table 2.** Comparison of C activation by Comirnaty and Doxebo in 75% human serum.

| Test agent | *sTCC* (sC5b-9, µg/mL)* |  | *CAI* | *SCA* | Zym | CMT/Dox<sup>##</sup> |
| --- | --- | --- | --- | --- | --- | --- |
|  | Mean | SEM (n=5) | (mg/mL)** | (*TCC/CAI*)*** | %<sup>#</sup> |  |
| Comirnaty | 2.3 | 0.6 | 1.0 | **2.3** | 0.5 | 4.6 |
| Doxebo | 3.1 | 1.1 | 6.4 | **0.5** | 0.1 |  |
| Zymosan | 148.4 | 22.9 | 0.5 | **494.8** | 100.0 |  |

Experiments like those shown in Figure 2 and 3, except that the 5 serum donors were different.
\**sTCC*, Mean ± S.E.M. of baseline corrected sC5b-9 readings after incubation with the test agents (Column 1) for 30 min; \*\*, *CAI*, concentration of C Activating Ingredient (mg lipid/mL), which is the sum of different lipid concentrations in Comirnaty and Doxebo, and the dry weigh/mL in zymosan; \*\*\*, *SCA*, Specific C Activation (*TCC/CAI*); <sup>#</sup>Zym %, *SCA* related to that of Zymosan (%); <sup>##</sup>Comirnaty (CMT) *SCA*/ Doxebo (Dox) *SCA* (ratio).

### Pathways and relative efficacies of C activation by Comirnaty and Doxebo

Beyond quantitating C activation by Comirnaty and Doxebo, **Figure 2** also gives information on the pathways of activation by the two nanoparticles. Namely, the finding that Doxebo caused significantly higher rises in C4d than Comirnaty (Figure 2C) indicates stronger involvement of the classical pathway in Doxebo-induced activation compared to Comirnaty, which difference lessened in the alternative pathway Bb assay (D). This attenuation of difference is consistent with alternative pathway amplification of classical pathway C activation, increasing the range of Bb changes to a 4-fold higher concentration level (see MWs in the legend).

The proposal of primarily alternative pathway activation by Comirnaty got strong support from the highly significant correlation between the vaccine-induced rises of C5a and both sC5b-9 and Bb (Figure 3A,B), but not between C5a and C4d (Figure 3C).

**Figure 3.**
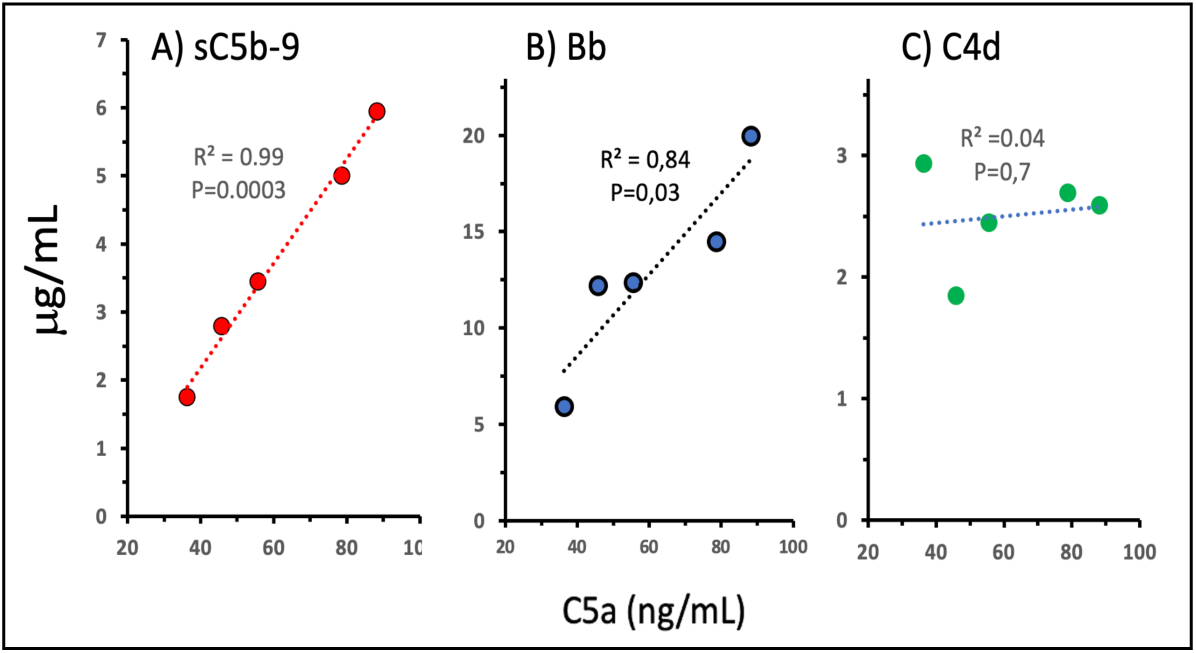
Correlations between C5a release and other reaction markers during Comirnaty-induced C activation in 75% human serum. These data were derived from the experiment also shown in Figure 2. As indicated by the R^2^ and P values near the regression lines, there were significant correlations between C5a and sC5b-9 (A) and C5a and Bb (C), but not between C5a and C4d (B).

Regarding the near equal rises of sC5b-9, C5a and Bb caused by Comirnaty and Doxebo (Figures 2A, B and D), although the lipid amount was ∼6-fold lower in Comirnaty compared to that in Doxebo, this result raised the possibility of substantial differences in C activating efficacy. Quantifying the latter by dividing the vaccine-induced C activation, expressed as maximal rise of sC5b-9 (*sTCC*), by the weight of assumed C activating ingredient (*CAI*), the resulting ratio “Specific C activation, *SCA*”, was 4.6-times higher in Comirnaty than Doxebo (Table 2).

### Effects of Spikevax and naked mRNAs

To further dissect the individual contributions of Comirnaty ingredients to C activation, we tested the C activating effects of Spikevax and naked mRNAs. Like Comirnaty, Spikevax is also an mRNA-LNP, but the ionizable and PEGylated lipids differ in the two vaccines (see Figure 4 legend). Thus, if C activation is due to a particular lipid, the two vaccines’ C activation is expected to be different. Likewise, if C activation is fully due to the lipid, its mRNA should not have any C activating effect. These presumptions were tested by incubating human sera with Comirnaty and Spikevax at equimolar lipid concentrations, as well as with vaccine mRNA-equivalent amounts of commercially obtained 5-methoxyuridine-modified SARS-CoV-2 mRNA.

**Figure 4.**
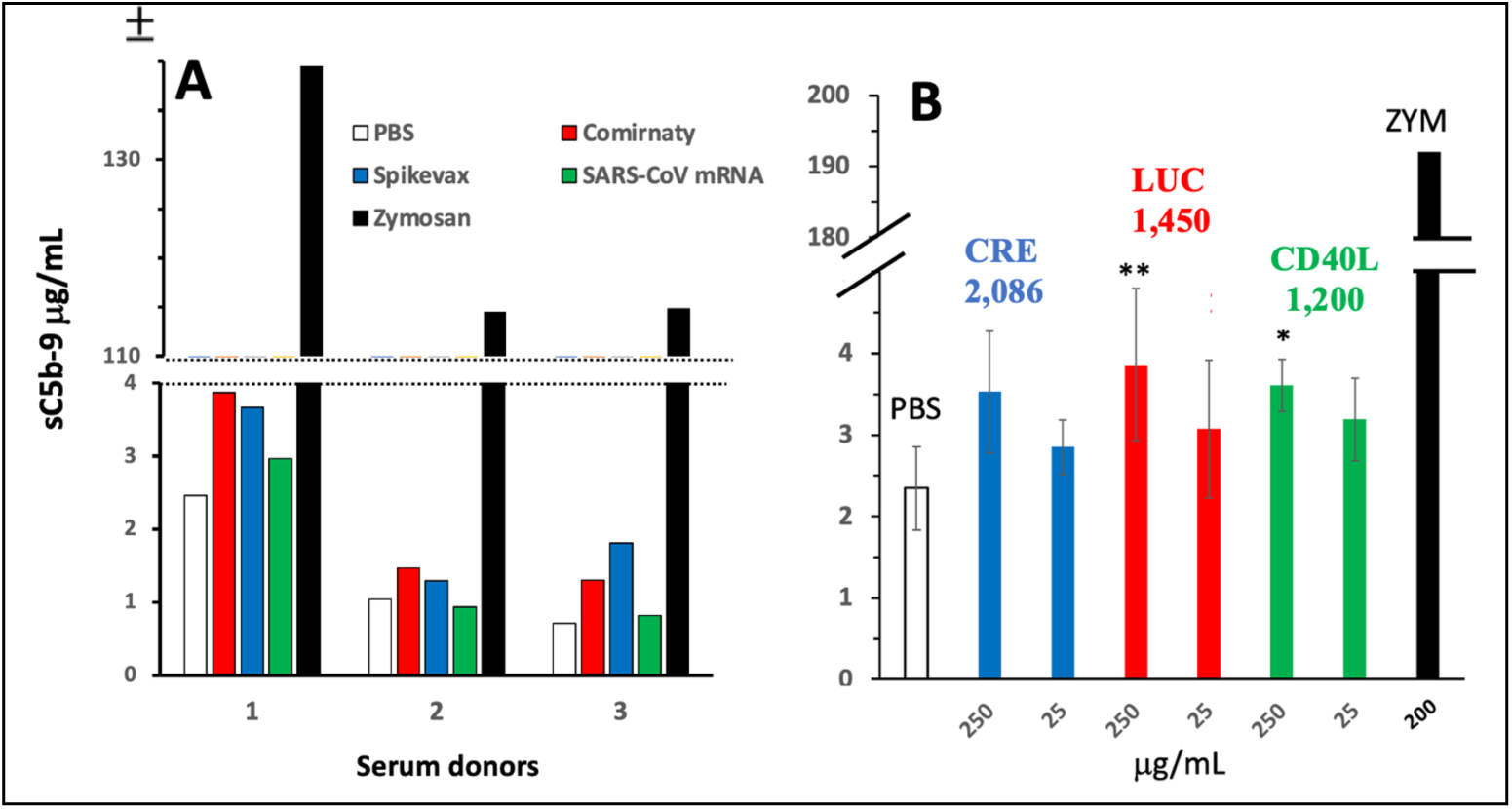
A). Complement activation by the two mRNA vaccines and SARS-CoV-2 mRNA (A) or synthetic mRNAs (B) in solution. An experiment like that in Figure 2, except the blood donors were different. In A, the final concentrations of Comirnaty and Spikevax were: 20 μg mRNA/mL (0.52 mg/mL total lipid) and SARS-CoV-2, 20 μg mRNA/mL. The lipids in Comirnaty are listedf in Table 1. Those in Spikevax; heptadecan-9-yl 8-((2-hydroxyethyl) (6-oxo-6-(undecyloxy)hexyl)amino) octanoate (SM-102), polyethylene glycol-2000-dimyristoyl glycerol (PEG-DMG), DSPC and cholesterol (total lipids: 3.86 mg mL^−1^). In B, the mRNAs tested were coding for the CD40L, Luciferase and_Cre-recombinase proteins, transcribed from the moCD40L, rstsFluc and Cre-recombinase genes, respectively. Their sequences are shown in STable 3-5 and nucleotide numbers on top of the bars in panel B. The abscissa shows the concentrations of these mRNAs added to serum: Zym, zymosan. Mean SEM, n= 5 different sera. *, **, significant differences compared to PBS at P<0.05 and P< 0.01 (ANOVA followed by Dunnetts’ post-hoc test).

As shown in Figure 4A, Comirnaty and Spikevax caused near equal low level C activation in 3 human sera, while SARS-CoV-2-mRNA had no or less effect. The Comirnaty-Spikevax near equivalence was confirmed in another measurement as well, performed as part of the inhibitor testing experiment presented below (see Figure 5B,C). Likewise, the lack of C activating effect of SARS-CoV-2 mRNA was reinforced in another experiment shown in STable 2. These data therefore suggest that it is the PEGylated lipid coating of mRNA-LNP nanoparticles that is responsible for C activation rather than a special feature of (a) lipid constituent of the vaccine, and that the vaccine mRNA has no major contribution to this process. Nevertheless, the latter conclusion had the limitation that the SARS-CoV-2 mRNA contains 7-times more nucleotides than the SP-coding vaccine mRNA (29,903 nt vs. 4,284)^61, 62^, and in lack of information on the size dependence of C activation by free mRNA, it is not *a priori* excluded that the smaller polyanion is a stronger C activator. Therefore, we engaged in testing 3 other, well-characterized mRNAs differing in nucleotide numbers, at near equal and a 100-fold higher concentration than the mRNA level is in the vaccine.

**Figure 5.**
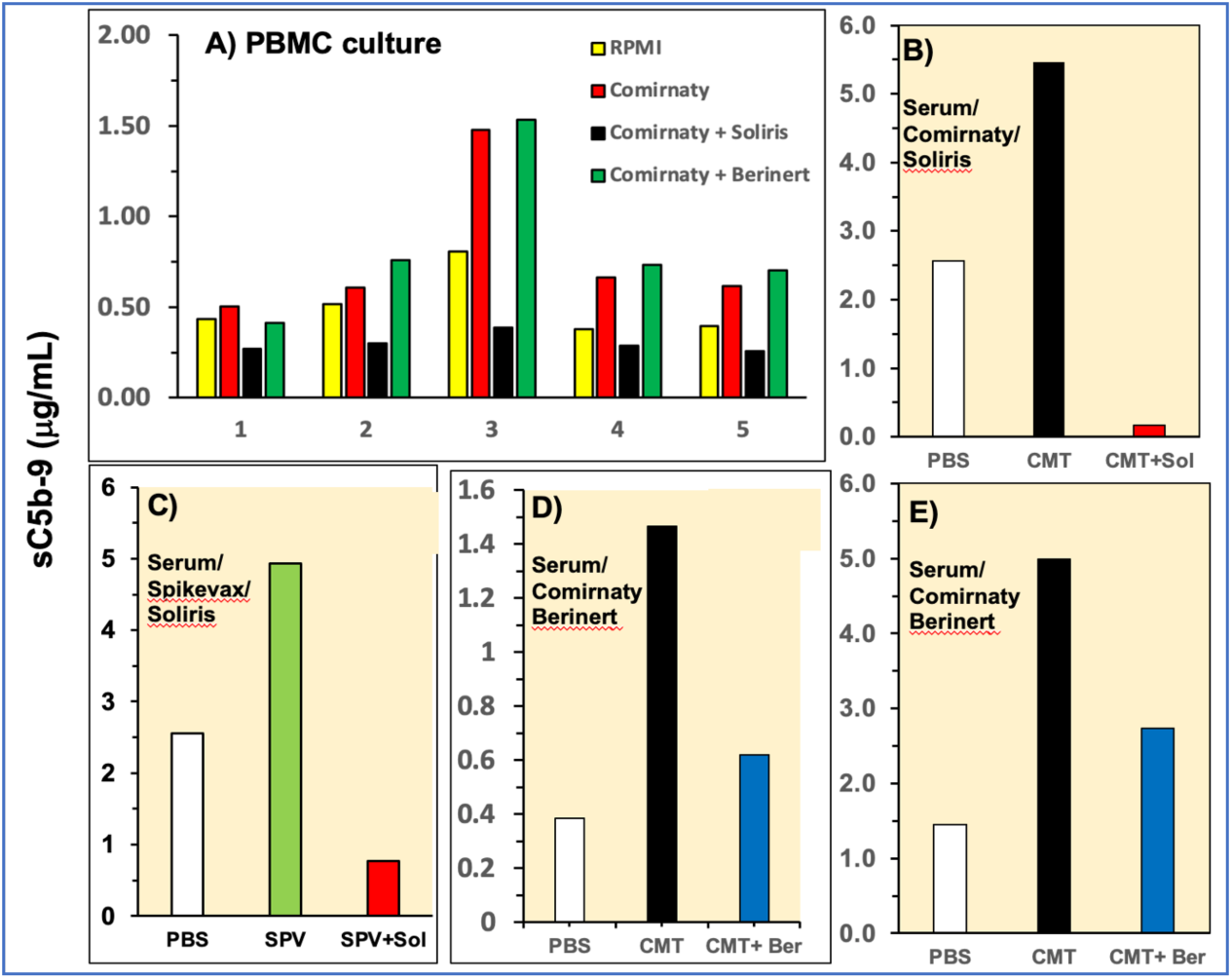
Inhibition of Comirnaty-induced C activation by Soliris in PBMC cultures from 5 subjects (A) and in an individual 75% sera (Figure 5B). Similar, individual measurements using Spikevax as C activator (Figure 5C), and, in Figures 5D and E, Comirnaty as activator and Berinert as inhibitor. Yellow bars in Figure 4A and empty bars in C-E show the 20% and 75% sera baselines, respectively.

As shown in Figure 4B, statistically significant rises of sC5b-9 in 2 of 3 mRNAs samples were seen only at the 100-fold higher mRNA level, but the differences were very small relative to the dose escalation. These data argue against an active role of mRNA in C activation by Comirnaty, however, as discussed in the Discussion section, the evidence of a lack of C activation by free mRNA may not be conclusive as the mRNA in the vaccine is complexed to lipids^62^.

### Effects of C inhibitors on C activation in 20 and 75% sera

Beyond featuring C activation by Comirnaty, a major goal of our study was to explore possibilities for inhibiting it. To this end we tested the efficacies of two clinically available C inhibitors, the C5 blocker Soliris and C1 inhibitor Berinert. Figure 5 A-E show two types of experiments iterating the test systems and different variables. In PBMC culture, Soliris, but not Berinert, exerted complete inhibition of sC5b-9 formation in all 5 sera (Figure 5A). This effect of Soliris was reproduced in 2 tests using 75% sera and either Comirnaty, or Spikevax as test agent (Figures 5B and C). Spikevax, Moderna’s mRNA vaccine, was tested to find out if the 2X higher mRNA concentration in a lipid-normalized sample had a major impact on C activation. The 2-fold increase that the lipid-normalized Spikevax caused in sC5b-9 (Figure 5C) and the inhibition of C activation by Soliris were identical to the changes seen with Comirnaty (B), confirming the causal role of vaccine lipids in C activation.

### Cytokine induction by Comirnaty in PBMC culture and the impacts of heat inactivation and C inhibitors

As shown in Figure 6A-F, Comirnaty induced significant production of IL-1α, IL-1β, IL-6, IL-8, IFN-γ, and TNF-α in the same 20% autologous serum-supplemented PBMC culture that were tested for C activation in Figure 1. In contrast, IL-2, IL-4, and IL-10 did not display any response to the vaccine (data not shown). The secreted amounts of responder cytokines increased in the following order: IL-1α < IFN-γ < IL-1β < TNF-α < IL-6 < IL-8. Among the latter changes, the rises of IL-1α, IL-1ϕ3 and TNF-α were eliminated using the heated serum supplement, while those of IL-6 and IFN-γ were entirely serum independent. The most abundantly produced IL-8, a chemokine that responds to most lipid and polymer based nanoparticles^63^, showed partial serum dependence, at least in 4 of 5 individuals.

**Figure 6.**
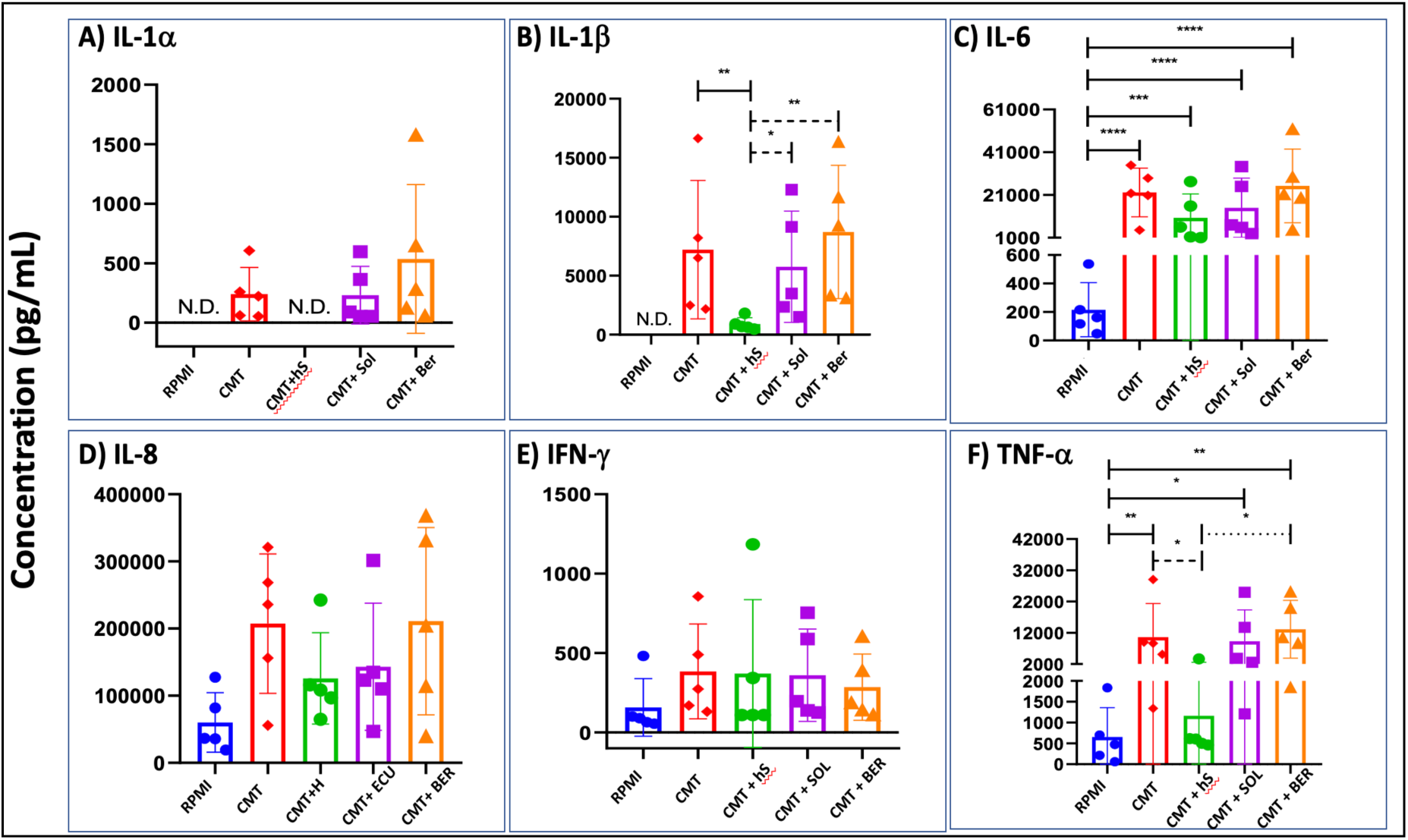
Cytokine production by Comirnaty in PBMC culture supplemented with 20% autologous serum with or without heat inactivation and C inhibitors. Differently colored symbols from 5 blood donors distinguish the treatments. The concentrations of Comirnaty, Soliris (SOL), and Berinert (BER) were 16 μg/ml (mRNA), 1 μM, and 1 μM, respectively. Bars show the mean along with the individual points. Statistical comparisons were done using one-way ANOVA followed by Tukey’s post hoc tests. *, **, ***, and **** mean p< 0.5, 0.1, 0.05, and 0.01, respectively. Other experimental conditions are described in the Methods. N.D., non-detectable

Since heat inactivation of serum supplement did not reduce the production of IL-6 (Figures 6C and E), the negative impact of heat inactivation is unlikely to be due to a universal toxicity on the cells. On the other hand, since C was activated by Comirnaty in all native-serum supplemented PBMC cultures, and since the heat-lability of C activation is well known, it seemed certain that the serum-dependence of cytokine release reflects a dependence on heat-labile C proteins. Nevertheless, we wished to get alternative evidence as well, namely the use of Soliris and Berinert which inhibited C activation in PBMC cultures (Figure 1) and serum (Figure 5) partly or fully. Thus, we measured Comirnaty-induced cytokine release in the presence of these drugs, but unexpectedly, they did not reduce the cytokine’s induction (Figure 6A-F).

In sum, just as in the C response, there is substantial variation in the presence and serum-dependence of different cytokines’ response to the vaccine. Among the cytokines that respond to the vaccine, some are fully serum dependent, some others not at all, and there are ones which are uncommitted in this regard. Whether it is a heat-sensitive C or another molecule in serum that is responsive for mediating the serum signal, remains to be established. If it is a C protein, the effect of heat cannot be reproduced by single C1 or C5-targeted inhibition of C activation.

## DISCUSSION

### Controversy over the severe adverse effects of mRNA vaccines

According to the most recent directives of the US Centers for Disease Control and Prevention and the World Health Organization, the mRNA COVID-19 vaccines are effective and safe^1–4^. This was first proven in the mentioned (see Introduction) multinational, placebo controled, double-blinded phase 2/3 trial of BNT162b2, conducted by Pfizer and BioNTech in 2020^7^. The study involved 21,621 vaccine recipients, among whom 20.7% reported some type of AE and 1.1% any kind of SAE. The corresponding numbers among 21,631 placebo recipients were 5.1% AE and 0.6% SAE, leading the authors to conclude that the “incidence of SAEs was low, similar in the vaccine and placebo groups”, and “the safety of the vaccine over a median of 2 months was similar to that of other viral vaccines”^7^. Of note, the frequency of SAEs in recipients of all other currently used vaccines is in the 10^−4^-10^−2^ % range^64–66^.

While the scientific^67, 68^ and top level (US senate, UK parlament) public debate on the safety of mRNA vaccines and causes of excess death are still going on, there is increasing number of scientific publications on the AEs of mRNA vaccines (Table 1)^8–41^. Yet, consensus on their implications has not been reached, and the mechanisms of unusual, post-COVID-like “special interest” SAEs^5, 6^ remain to be understood.

### Study rationale

C activation, a phylogenetically ancient, basic mechanism involved both in innate and adaptive immune responses^69^, is a proinflammatory process that can be a trigger, a consequence or an epiphenomenon of proinflammatory cytokine release by immune cells, another essential process feeding into inflammations. Differentiating among these relations in the case of Comirnaty may help to assess the use of C inhibitors against the inflammatory AEs of the vaccine. In particular, in case of causal relationship, C inhibition could be effective against both processes. For this reason, the main goal of this study was the analysis of vaccine-induced C activation and exploration of its possible role in cytokine production.

The present experiments extended previous results in different models providing evidence for C activation by Comirnaty^17^ and cytokine induction by PEGylated LNPs similar to the vaccine^70–72^. As a clinically relevant in vitro system to study immune cell responses, such as cytokine release^46, 73–75^, we used PBMC cultures supplemented with native, instead of heat-inactivated serum; a deviation from tissue culturing standard porocedures enabling the assessment of the role of C activation in the observed immune cell changes^46^. Doxebo was used as control for Comirnaty because of the similaries of their nanostuctures^62^ and wealth of information on C-activation and related hypersensitivity/anaphylactoid reactions caused by Doxebo’ parent drug, Doxil^44, 76–78^. Finally, zymosan, the gold standard of C activation was used as positive control, as in all our previous in vitro^17, 46^ and in vivo^17, 79, 80^ studies on CARPA.

### C activation by Comirnaty: old and new information

Complement activation by liposomes and other therapeutic or diagnostic nanoparticles has been known for decades^81^, but the fact that mRNA-LNPs can also cause it has generally been overlooked in the mainstream literature on COVID-19 vaccines. We have previously reported C activation by Comirnaty in pig serum in vitro^17^ and in pig blood in vivo, after i.v. injection of the vaccine in anti-PEG hyperimmune pigs^19^. The present reproduction of the phenomenon in human serum implies that it is not species-specific. To our best knowledge, the proposal that C activation by the vaccine and PEGylated liposomes are qualitatively similar, and is likely due to an intrinsic property of PEGylated nanoparticles, is also original. Yet further new informations in our study include the feasibility of using native serum-substituted PBMC cultures to measure immune cell functions, the accumulation of sC5b-9 upon heat treatment of serum and the absence of biologically relevant C activation by naked mRNAs.

### The mechanism of complement activation by Comirnaty and Doxebo

In theory, each constituents of Comirtnaty and Spikevax can activate C, namely the phospholipids^82^, the PEGylated lipids^16, 19, 83^, the cholesterol, at high membrane concentration ^84, 85^, the nucleic acids^86^, and even free PEG, in solution^87^.

Looking closer at the above possibilities, the fact that Comirnaty and Doxebo caused the liberation of the same C clevage products points to what is common in these nanoparticles, namely their size, shape and the presence of PEGylated lipid membrane coat^62^. The lack of C activation by naked mRNAs strenghtens the idea that C activation was due mainly to the lipids, most likely because they form membranes providing sufficiently large platform for antibody binding, C3b deposition and assembly of C3 and C5 convertases. However, the percentage of membrane-forming phospholipids in Comirnaty was ∼4 times less compared to Doxebo, and comparing only the PEGylated lipid percentages, the ratio was 3-fold lower in Comirnaty (STable 1). These ratios taken together with the ∼6 times lower total lipid concentration in the Comirnaty samples (Table 1, Figure 2) and ∼5 times higher specific C activation of Comirnaty vs. Doxebo (Table 2) raises the possibilty that structural differences between these nanoparticles that exist beside the similarities, entail a major amplification of C activation by the vaccine through an unclear mechanism.

A possible solution for the above paradox is offered by a recent study, showing that unlike Doxebo, Comirnaty consists of soft, fragile nanoparticles prone for disintegration in water to yield lipid fragments and snake-like twisty nanostructures, tentatively identified as mRNA lipoplexes^62^. The breakup of vaccine nanoparticles obviously results in the expansion of serum-exposed surfaces, thus platform for additional C activation. In addition, the possible emergence of mRNA-lipoplexes also increases the chance of reaction triggering, as complexation of nucleic acids with positively charged polymers were shown to be strong C activators^86^. Finally, it should be mentioned that the common phospholipid in mRNA-vaccines, DSPC, significantly increased C activation by liposomes^84^, and that ALC-0315 in Comirnaty is an unnatural lipid, whose clusters^62^ might trigger C reaction directly, or after association to lipoproteins, as mentioned above^88, 89^.

As to the role of PEGylated lipids, we found that unlike Comirnaty, Doxebo caused a significant rise in C4d, indicating classical or lectin pathway activation. This observation is in keeping with previous data on C-mediated anaphylactoid reactions to Doxil or Doxebo, triggered by the binding of anti-PEG antibodies to PEG on the vesicles^18, 19, 70, 83, 90, 91^. Although Comirnaty also contains PEG, its surface density is lower than that in Doxebo (1.5% vs 5%), which may explain the small but consistent C4d rise in 3 of 5 sera in Figure 2C. In other experimental systems, e.g., anti-PEG hyperimmune pigs^19, 83^, or even humans with extremely high anti-PEG antibody levels in their blood^18^, the involvement of classical pathway in C activation by PEGylated nanoparticles may become more prominent.

The mechanism of alternative pathway activation by both Comirnaty and Doxebo is less clear. The mentioned instability of Comirnaty in water^62^ raises the possibility of rapid distribution of its main lipid, ALC-0315, to lipoproteins in plasma, entailing direct binding of C3, in a way similar to that observed with Cremophor EL^88, 89^. In addition, what explains the alternative activation by both NPs, is acceleration of classical pathway C activation via alternative pathway amplification^92^. In support of this explanation, conversions of the average Comirnaty-induced rise of C4d (∼1 ug/mL, Figure 2B) and Bb (∼10 μg/mL, Figure 2D) to molar ratios (see MWs in the legend to Figure 2) gave ∼7-fold more Bb molecules than C4d, which is consistent with the amplification of C activation by the alternative pathway positive feedback loop^92^.

### The relationship between Comirnaty-induced complement activation and inflammatory cytokine production

It is well known that the anaphylatoxins C3a and C5a, and the proinflammatory cytokines, including IL-1β, IL-6, TNFα and IF-γ, are crucial mediators of inflammation. A crosstalk between these two arms of innate immunity has been described in different systems^46, 79, 93–95^ but not for mRNA-LNP vaccine-induced AEs. In theory, Comirnaty-indued C activation can enhance cytokine release via anaphylatoxin receptors and via opsonization of nanoparticles, thus enhancing their uptake and binding to inflammation-signaling pattern (and/or danger) receptors of immune cells. In support of the latter possibility, previous studies from our laboratory showed temporal correlation between C activation and release of inflammatory cytokines by zymosan in pigs^79^, and suppression of zymosan-induced IL-6 release in PBMC culture by factor H, an inhibitor of altrnative pathway C activation^46^. As for the observation that heat-treatment of serum suppressed the production of IL-1α, IL-1β and TNFα, the phenomenon is most likely due to inactivation of (a) heat labile C protein(s) that play (a) causal role(s) in the secretion of the above and possibly other cytokines. However, development of anti-complementary activity due to IgG aggregation and/or inhibition of phagocytic uptake of vaccine NPs cannot be excluded^55, 96^, either.

We have tested the effects of of Soliris and Berinert, two clinically applied C inhibitors, and while they inhibited C activation, they had no inhibitory effects on the production of either studied cytokine. Although we have not further analyzed the possible reasons for these negative results, one explanation may be that C-dependent cytokine release is mediated by simultaneous actions of anaphylatoxins and other activation byproducts, and targeted suppression of C activation via the classical pathway by Berinert, or C5 conversion by Soliris, may not be effective in suppressing cytokine release when applied individually. This implies that the C system needs to be simultaneously inhibited at several check points to achieve cytokine suppression, for which many options exists^97–103^. The clinical efficacy of Soliris in attenuating cytokine storm in severe COVID-19^104–107^ provide evidence that C inhibition may be useful against cytokline release, requiring further studies to explore the preconditions of this beneficial effect.

### Clinical implications of vaccine-induced C activation

According to the prescription information of PEGylated liposomal doxorubicin (Doxil)^108^, the iv. administered drug can cause acute infusion reactions in up to 11% of patients, and these reactions can be “serious, life-threatening, and fatal”^108^. Human^77^ and animal studies^44, 78, 109^ have previously linked these HSRs to C activation by Doxil, while other studies have shown that Doxil and Doxebo are near equivalent in causing C activation^44^. Thus, despite the low level of C activation in vitro and low number of studied sera, the consistent finding of C activation by Comirnaty supports the implication of C activation in Comirnaty-induced HSRs and anaphylaxis^16, 17, 19^. The latter was listed in Comirnaty’s postmarketing survey, updated in Oct 2023, as the 3rd most frequent AE after the cardiac and gastrointestinal complications^110^. Although the i.m. administration of the vaccine is not comparable with the infusion of Doxil, the 5-times greater C activating potency of the vaccine and its administration as i.m. bolus could trigger similar C chain reaction as triggered by the few mg Doxil lipid that reaches the blood during the first few minutes of very slow infusion. The rise of HSR then depends on the sensitivity of the vaccine recipient to allergic reactions, as association of HSRs to Comirnaty and allergic constitution is well reflected by the exclusion of severely allergic, anaphylaxis-prone individuals from vaccination with mRNA vaccines.

As to the low level of in vitro C activation by Comirnaty and Doxebo in comparison with the effect of zymosan, it does not necessarily imply low biological activity in vivo. Anaphylatoxins are among the strongest proinflammatory substances in the body^111–114^, C5a being the most effective chemoattractant for neutrophil leukocytes. In a previous study, we injected 330 ng/kg rhuC5a to pigs, which raised the level of C5a in their blood to ∼40 ng/mL and caused detectable hemodynamic changes as a manifestation of C activation-related pseudoallergy (CARPA)^115^. This biologically active concentration is within the range of C5a rise in human serum caused by Comirnaty and the liposomes in the present study (Figure 2B). Yet in another previous animal study, i.v. injection of Comirnaty in anti-PEG antibody hyperimmune pigs at a dose containing ∼0.3 μg/kg mRNA (theoretically resulting ∼10 ng/mL mRNA in plasma) caused lethal anaphylactic shock in 6 of 6 pigs^19^. This plasma level is ∼600-fold lower than the mRNA level in the present in vitro study causing major elevation of C5a (Figure 2B).

Regarding the question, what is the threshold of C activation biomarker elevation leading to clinical symptoms, a study in Doxil-treated cancer patients showed increased susceptibility for CARPA at >2-3-fold drug-induced rise of plasma sC5b-9 levels^77^, a rise that was reached in 4 of 5 serum in the present study (Figure 2A). In sum, several lines of indirect evidence point to a role of C activation in the HSRs and anaphylaxis to Comirnaty, and the speculation below gives reason to propose that C activation may underlye the subacute and chronic AEs of Comirnaty, as well.

### Link between proinflammatory impact and vaccine-induced SAEs: a hypothesis

By exposing a connection between the release of proinflammatory cytokines and C activation, our current study raises the possibility that the inflammatory side effects are due, at least in part, to additive or synergistic proinflammatory actions of anaphylatoxins and inflammatory cytokines, particularly if these innate immune stimulations recur in the same subject. However, this proposal is realistic only if the different subclinical incubation times between the reaction trigger (i.e. anaphylatoxin/cytokine double hit) and start of symptoms can be reconciled. Examples for acute, subacute and chronic SAEs include the HSRs or anaphylaxis within minutes to hours^16–19^, heart inflammation^8–15^, coagulation disorders^21–24^, or skin diseases^25–28^ can occur within days to months, and autoimmune phenomena^20^ may arise without currently foreseeable time limit. A hypothesis that may explain these different latency periods postulates that the actions of effector immune cells, including the mast cells, basophil and neutrophil granulocytes, lymphocytes, macrophages and platelets, are discontinuous, graded in the sense that they manifest only after the activation state of their intracellular inflammatory signaling network reaches the reaction threshold. This threshold may be different in different cells, and each proinflammatory “hit” raises the activity state of effector cells to elevated, yet subclinical “alertness” stages until they add up and reach the reaction threshold. Variable activations of different arms of the multi-controlled intracellular inflammation-signaling network in these effector cells may explain the versatility of immune responses, whose entanglement remains to be a research challenge for the future.

In the case of COVID-19 mRNA vaccines, a potential extra “hit” after the initial response to anaphylatoxin and cytokine double hits may be C activation by the SP, since it is a potent C activator via the lectin pathway after binding to mannose-binding lectin, ficolin-2 and/or collectin-11^50^. In fact, there are studies showing the circulation of SP-expressing exosomes^116^ or the presence of the vaccine mRNA in a variety of tissues for weeks or months after mRNA vaccinations^8, 43, 117–120^, with potentials for SP translation and surface expression. A strong support for the above speculations is the globally increased occurrence of all AEs and SAEs after repeated vaccinations^110^ possibly because each mRNA vaccine booster may increase the risk of C activation due to the binding of a new wave of neutralizing anti-SP antibodies to SP expressing tissue cells^8, 43, 117–120^ or circulating exosomes^116^. As is well known, C activation is a pivotal function of antibody binding to different antigens, which may involve, in addition to IgM, various IgG subclasses depending on the conditions^121^. Such antibody-mediated C activation may join the intrinsic C activating activity of the SP via the lectin pathway^50^.

### Outlook

The creation of the mRNA-LNP vaccines was possible partly because nucleoside modification prevents the mRNA’s innate immune stimulating effect through Toll-like receptors^122^, and partly because the nucleoside modified mRNA could be delivered to immune cells after entrapment into LNPs^123^. It could not be anticipated that the vaccine can cause more or less increase in the prevalence of SAEs. As discussed, the phenomenon may involve inflammatory reactions due to innate immune stimulation by the vaccine, and/or delivery of vaccine mRNA to bystander (non-immune) host cells entailing potential expression of the SP antigen^43, 124, 125^ with subsequent aberrant immune responses, including autoimmune attack.

It should be emphasized that despite all above clinical information on rare adverse reactions and logics of mechanisms, the use of mRNA vaccines is justified until the favorable benefit-to-risk ratio validates it. The success of these vaccines has already resulted in significant progress in using the LNP technology for developing new gene therapies against a variety of diseases^123, 126–130^. Therefore, at this stage, the clinical significance of our findings is more in the future than the present, laying the groundwork for concepts that may guide the elimination of inflammatory SAEs to PEGylated LNPs. Combination pharmacotherapy with C inhibitors may offer a promising solution to this vexing problem.

## Author contributions

Conceptualization: BT, GS, JS; Experimentation: BT, TM., LD, GS, RF, PB; Data curation: HF, SK, RS, KAG, TR; BT, TM., LD, GS, RF, PB; Formal analysis: TM, GTK, SK; Writing: JS, GS, TB; review & editing: BT, GS, KAG, TM, RF, TR, PB. Supervision: GS, JS, RS, TR.

## Data availability

Data will be made available on request.

## Acknowledgments

“Sincere thanks are due to Drs Marina Dobrovolskaia (NCL, NCI, NIH, Frederick, USA) and Joshua Jackman (Sungkyunkwan University, Suwon, Korea) for their advice on the manuscript, and Mr. Zoltan and Miklos Szebeni for the grammar corrections. JS thanks the support of the Applied Materials and Nanotechnology Center of Excellence, Miskolc University, Miskolc, Hungary.

---

**STable 1.**
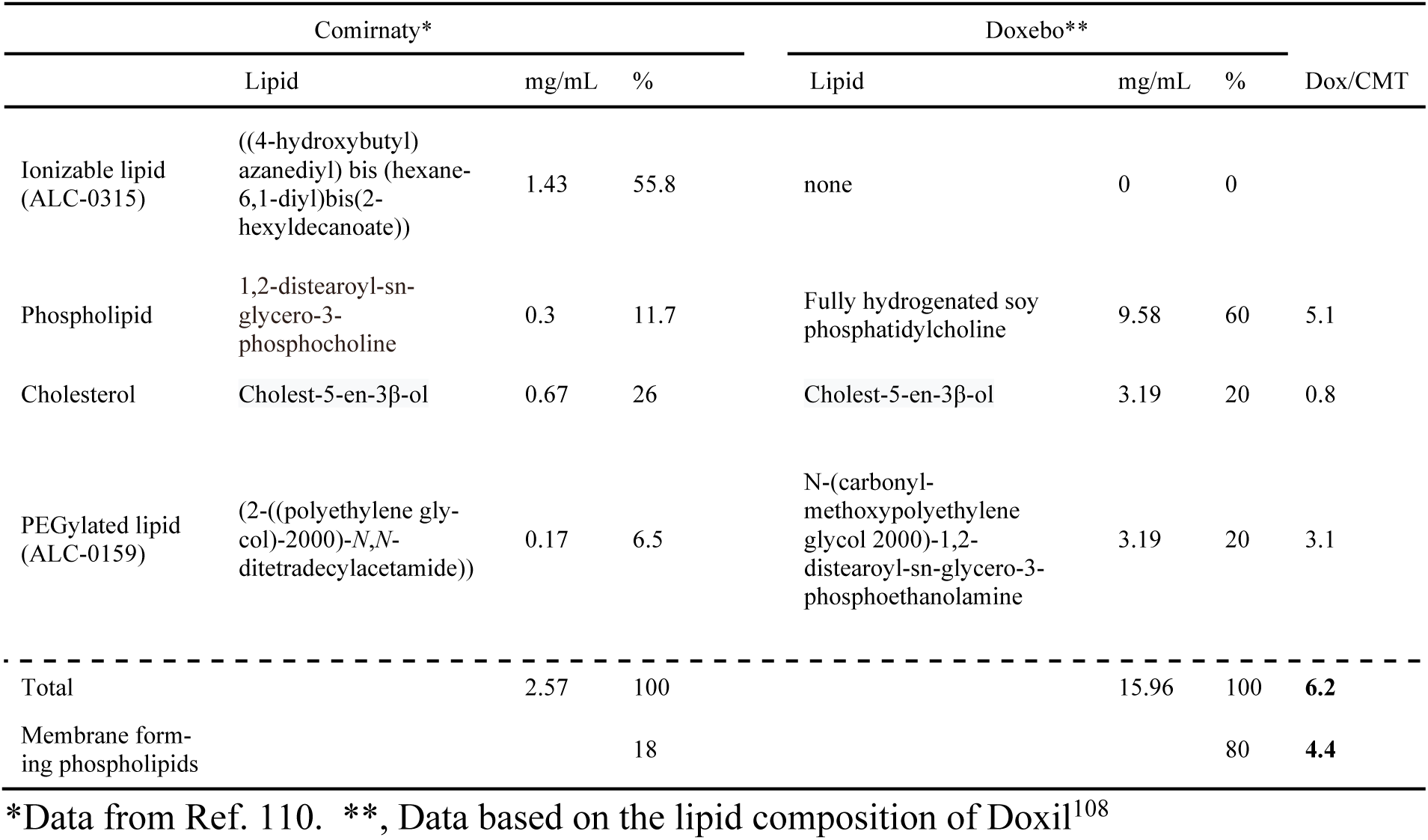
Lipid composition of Comirnaty and Doxebo.

| Comirnaty* |  |  |  | Doxebo** |  |  |  |
| --- | --- | --- | --- | --- | --- | --- | --- |
|  | Lipid | mg/mL | % | Lipid | mg/mL | % | Dox/CMT |
| Ionizable lipid (ALC-0315) | ((4-hydroxybutyl) azanediyl) bis (hexane-6,1-diyl)bis(2-hexyldecanoate)) | 1.43 | 55.8 | none | 0 | 0 |  |
| Phospholipid | 1,2-distearoyl-sn-glycero-3-phosphocholine | 0.3 | 11.7 | Fully hydrogenated soy phosphatidylcholine | 9.58 | 60 | 5.1 |
| Cholesterol | Cholest-5-en-3β-ol | 0.67 | 26 | Cholest-5-en-3β-ol | 3.19 | 20 | 0.8 |
| PEGylated lipid (ALC-0159) | (2-((polyethylene glycol)-2000)-N,N-ditetradecylacetamide)) | 0.17 | 6.5 | N-(carbonyl-methoxypolyethylene glycol 2000)-1,2-distearoyl-sn-glycero-3-phosphoethanolamine | 3.19 | 20 | 3.1 |
| Total |  | 2.57 | 100 |  | 15.96 | 100 | **6.2** |
| Membrane forming phospholipids |  |  | 18 |  |  | 80 | **4.4** |

\*Data from Ref. 110. \*\*, Data based on the lipid composition of Doxil<sup>108</sup>

**STable 2.** SARS-CoV-2 mRNA does not activate human complement.

|  | C4d (µg/ml) | Bb (µg/ml) | sC5b-9 (µg/ml) |
| --- | --- | --- | --- |
| PBS | 2.6 | 10 | 2.6 |
| Comirnaty | 5.1 | 17.2 | 5.5 |
| SARS-CoV mRNA* | 2.7 | 9.6 | 1.7 |
| Zymosan | 8.6 | 47.1 | 142.2 |

**STable 3.**
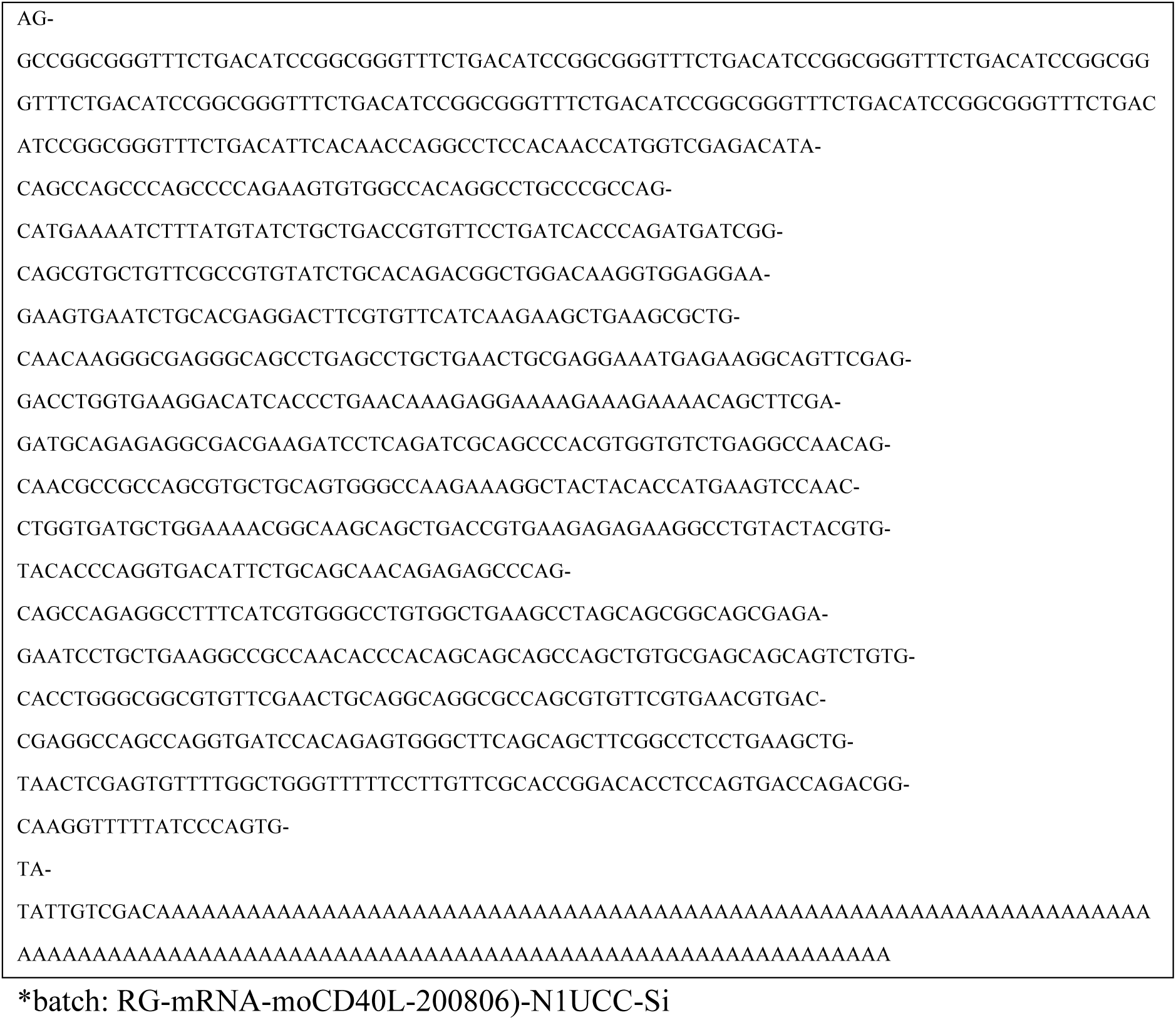
CD40L (moCD40L gene) mRNA sequence, 1,200 nt.

**STable 4.**
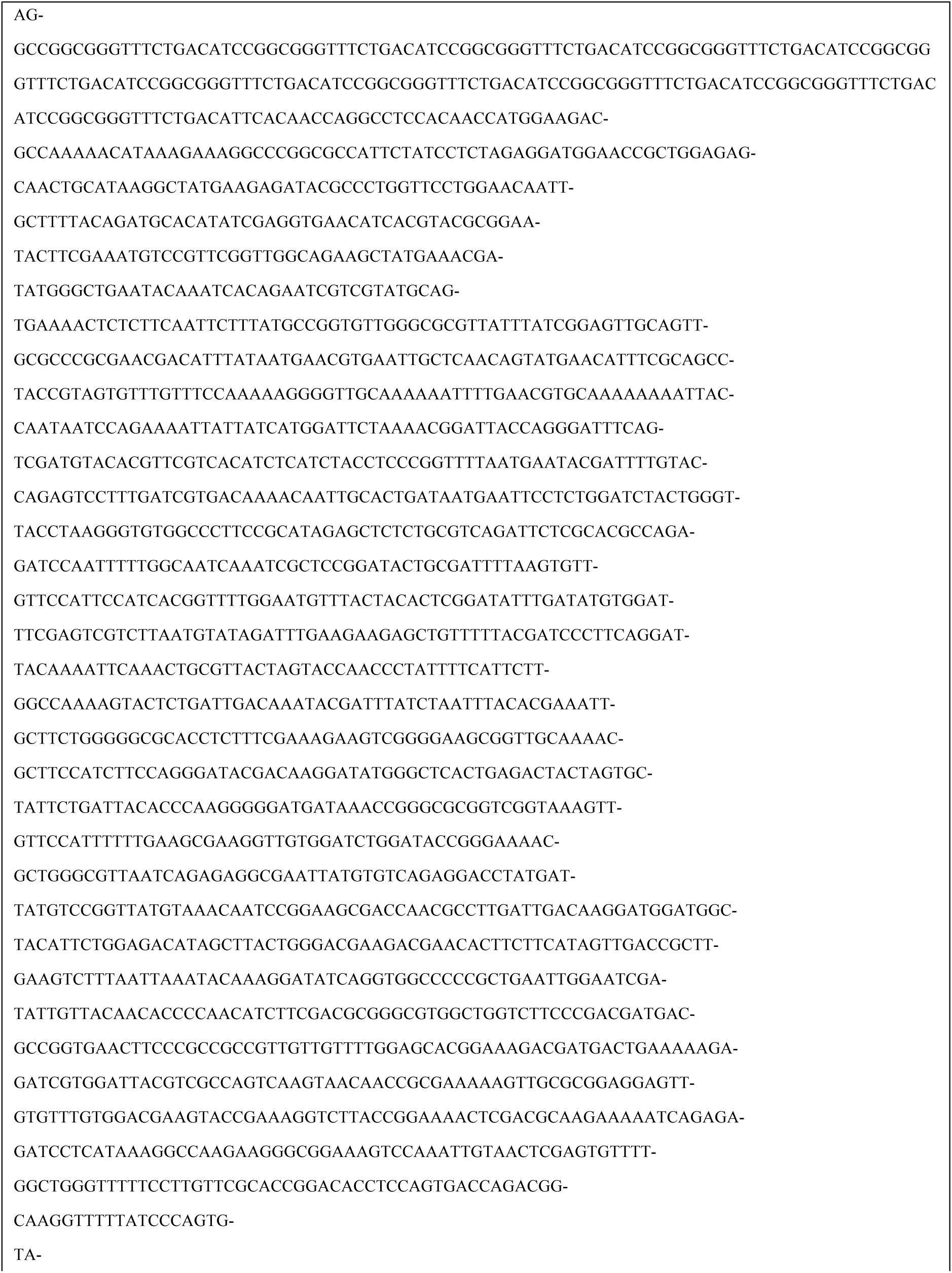
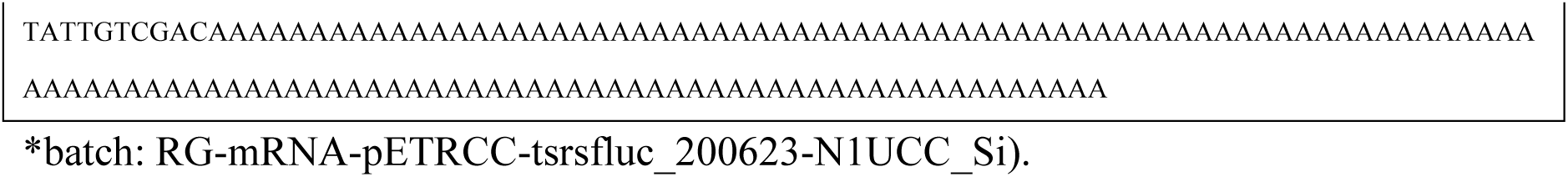
Luciferase (rstsFluc gene) mRNA sequence (1,450 nt)

**STable 5.**
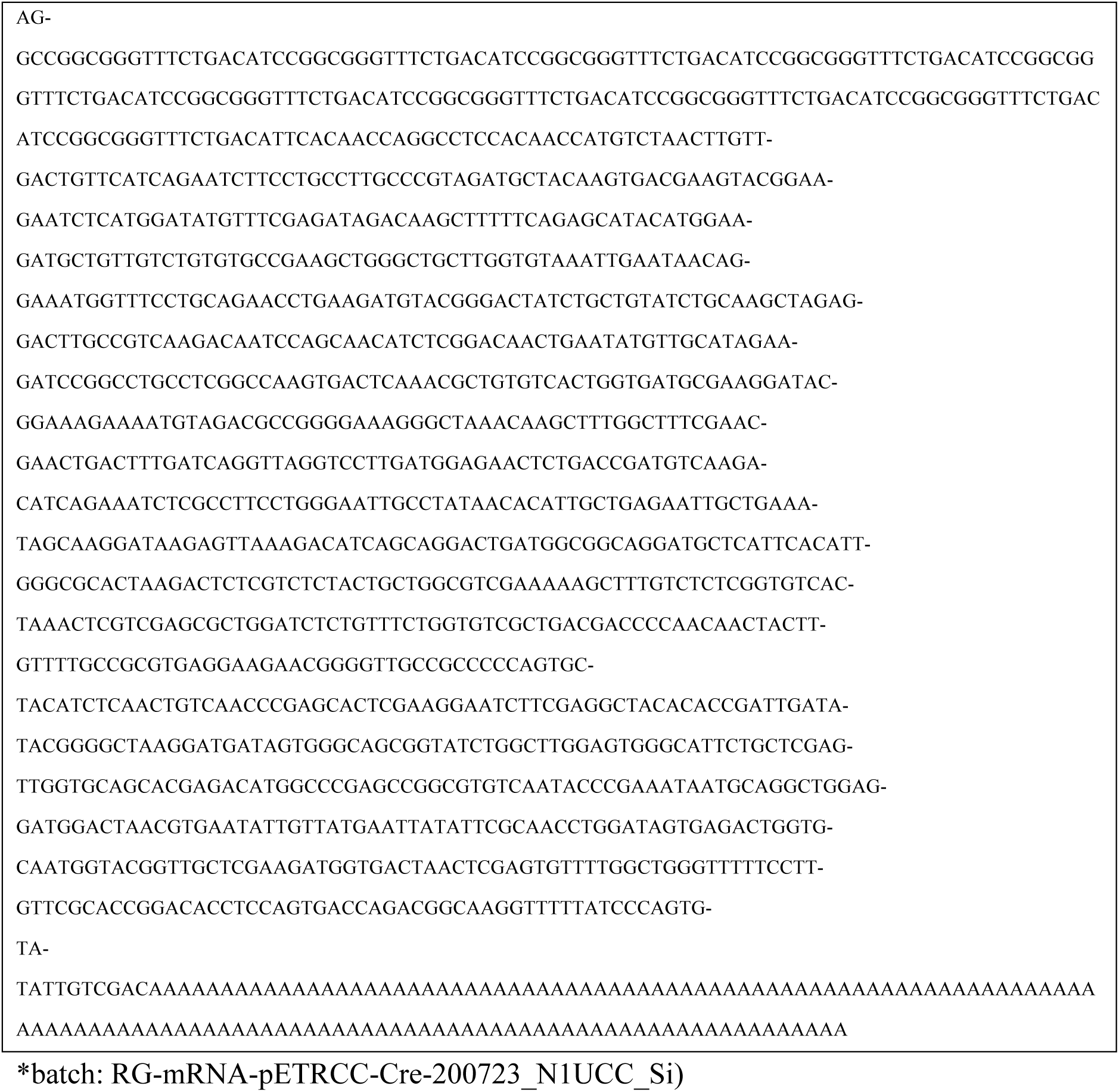
Cre-recombinase (Cre) gene mRNA sequence (2,086 nt)*

